# Pre-trial predictors of conflict response efficacy in human dorsolateral prefrontal cortex

**DOI:** 10.1101/2021.07.07.451322

**Authors:** Alexander B. Herman, Elliot H. Smith, Catherine A. Schevon, Mark Yates, Guy M. McKhann, Matthew Botvinick, Benjamin Y. Hayden, Sameer Anil Sheth

## Abstract

The ability to perform motor actions depends, in part, on the brain’s initial state, that is, the ensemble firing rate pattern prior to the initiation of action. We hypothesized that the same principle would apply to cognitive functions as well. To test this idea, we examined a unique set of single unit data collected in human dorsolateral prefrontal (dlPFC) cortex. Data were collected in a conflict task that interleaves Simon (motor-type) and Eriksen (flanker-type) conflict trials. In dlPFC, variability in pre-trial firing rate predicted the ability to resolve conflict, as inferred from reaction times. Ensemble patterns that predicted faster Simon reaction times overlapped slightly with those predicting Erikson performance, indicating that the two conflict types are associated with near-orthogonal initial states, and suggesting that there is a weak abstract or amodal conflict preparatory state in this region. These codes became fully orthogonalized in the response state. We interpret these results in light of the initial state hypothesis, arguing that the firing patterns in dlPFC immediately preceding the start of a task predispose it for the efficient implementation of cognitive action.

## INTRODUCTION

The ability to respond effectively to a stimulus can depend on the state of the brain even before the stimulus appears ^1–3^. In other words, our responses are determined not only by the neural activity driven by the response-driving stimulus, but by the way that activity interacts with ongoing neural activity ^4^. In the motor system, one expression of this idea is the *initial state hypothesis*, which holds that motor control involves a series of dynamical states and that initiation of motor control requires a particular state ^5–10^. Variability in performance, typically assessed with reaction times, corresponds in part to variability in pre-trial firing rates because those reflect the response of the system relative to the optimal response-driving initial state. We and others have proposed that dynamical principles relevant to the motor system may apply to non-motor processes, including higher level cognitive processes ^11–15^. We hypothesized, therefore, that the ability to implement a cognitive process may likewise depend on the initial state of the system.

We are particularly interested in conflict detection and resolution, a pair of complementary and relatively well-studied cognitive behaviors whose neuronal basis is beginning to be understood ^16–21^. Conflict typically refers to a competition between possible stimuli for attention and/or action and generally evokes slower reaction times, increased error rates, and disengagement from alternative tasks ^22,23^. This refocusing of mental resources is the basis of conflict resolution and presumably occurs in response to an internal detection of conflict and generation and propagation of a conflict signal. We hypothesized that the ability to deal effectively with conflict depends in part on variability in neural processes in conflict-relevant brain regions before the appearance of the conflicting stimuli.

The dorsolateral prefrontal cortex (dlPFC) is among the most studied brain regions for cognitive control, along with anterior cingulate cortex ^16,20,24-26^. dlPFC shows systematic changes in hemodynamic response, local field potential (LFP), and firing rate in the face of conflict (ibid.). We have recently proposed that these two regions play somewhat distinct, albeit complementary roles in conflict detection and resolution ^16^. We proposed that dlPFC is more associated with implementation, and thus potentially a closer cognitive analogue to motor areas (see also ^25–27^).

Here, we examined firing rates of single neurons in dlPFC while humans performed the multi-source interference task (MSIT), a task that manipulates two different forms of conflict, a motor (Simon) type and a perceptual (Eriksen flanker) type ^16,19^. We found that activity patterns in the period preceding the start of the trial predicted reaction time in dlPFC, even after regressing out prior trial reaction time and conflict type. At the individual cell level, we found evidence for a neural code for response times specific to each conflict type. These codes overlapped slightly but significantly at the population level, indicating the presence of a weak shared conflict-amodal code. These results endorse the idea that conflict resolution reflects the interaction between stimulus-driven activity and ongoing fluctuations in pre-trial activity, support the initial state hypothesis for cognitive actions, and suggests a mechanism by which the brain can respond both flexibly and efficiently to different conflict conditions.

## RESULTS

### Behavior

We examined responses of single neurons recorded in dlPFC in 9 human patients (**Figure 1A**). Task-related responses in this dataset were described in an earlier study, but pre-trial responses, the focus of the present study, were not analyzed ^16,28^. Participants performed the multi-source interference task (MSIT), a task that involves two independently manipulated types of conflict (**Figure 1B**).

**Figure 1.**
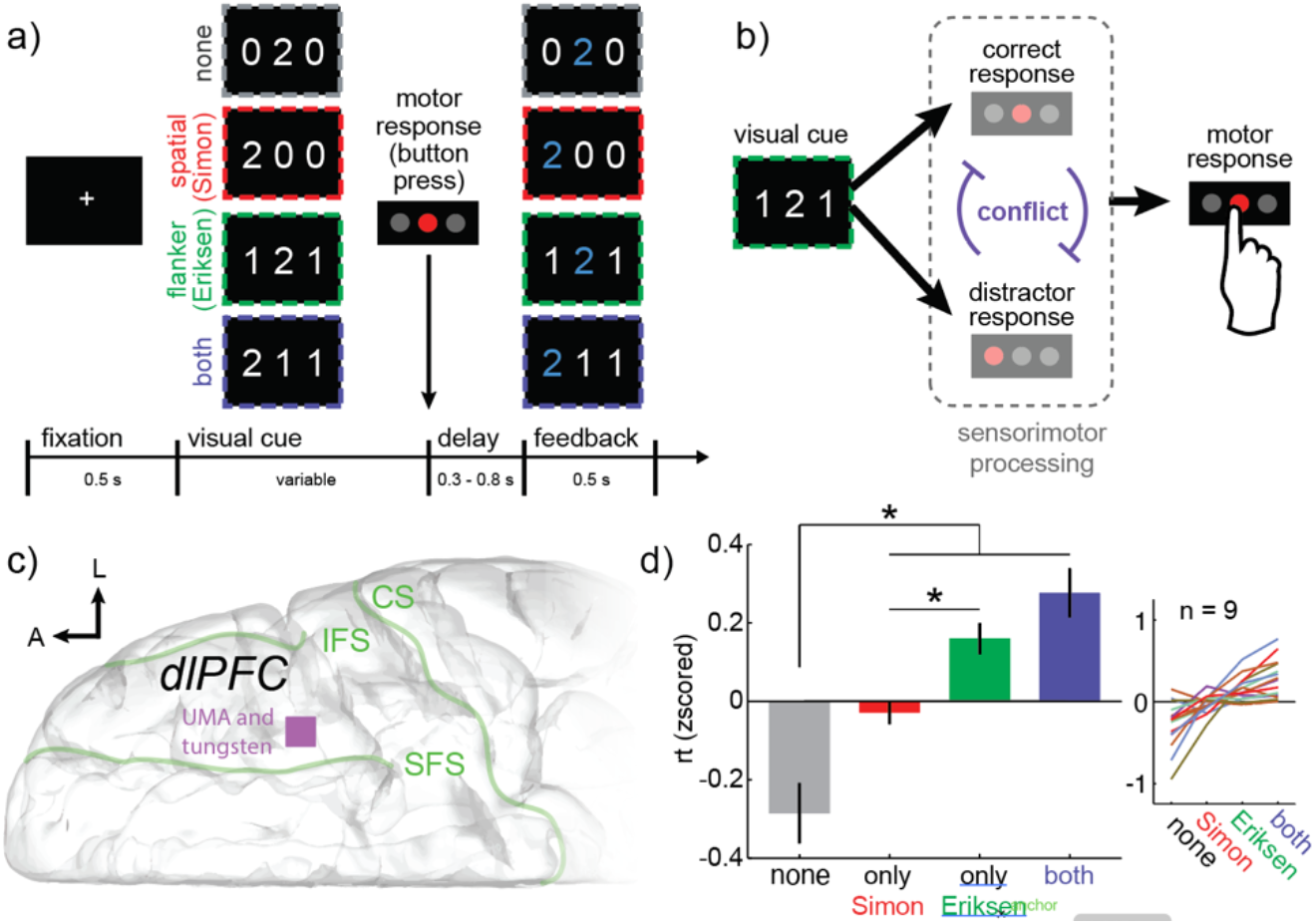
Multi-source interference task (MSIT) design, recording locations, and behavioral results. (A) Basic task design. Participants fixate on a central cross and then see a visual cue consisting of three numbers and has to identify the unique number with a button push. “correct response” is the left button if the target is 1, middle if 2, right if 3. Four example cues are shown here, and in each case, the target is “2” and the middle button is the correct response. This is most obvious for the first cue (“none”), where there is no conflicting information. In the other three examples, conflicting information makes the task more difficult. First, incongruence between the location of the target number in the 3-digit sequence and location of the correct button in the 3-button pad produces spatial (Simon) conflict (orange). Second, the distracting presence of numbers that are valid button choices (“1”, “2”, “3”) produces flanker (Eriksen) conflict (green). Trials can also simultaneously have both types (blue). B. The visual cues are associated with one or more sensorimotor responses. Every cue has a correct response, meaning the button press that corresponds to the unique target. Cues can also have one or more distractor responses, meaning the button press that corresponds to task-irrelevant spatial information (Simon) or flanking distractors (Eriksen). If and only if the correct response and distractor response do not match, then the cue causes conflict because only one button response can ultimately be chosen. C. Diagram of the intracranial implant showing the UMA and tungsten microelectrode recoding locations schematized as a purple square on the surface of dlPFC. sulcus. D. The average (mean) response times across subjects in each of the four task conditions and (right) the mean response times within each subject. Bars = standard error across subjects.

The validity of this task as a manipulation of conflict has been demonstrated ^16,19,29,30^. We therefore only briefly summarize the evidence that the task manipulates conflict. Most importantly, median reaction time in the Simon trials (1.5 sec) was significantly slower than no conflict trials (1.26 sec, p=0.017, z=2.402, ranksum=8438). Likewise, reaction times in the Eriksen trials (1.63 sec) was slower than in no conflict trials (p<0.001, z=4.51, ranksum=259). Finally, reaction times on both-conflict trials (1.71 sec) were longer than on Simon trials (p=0.006, z=2.71, ranksum=11428) although not compared to Eriksen: p=0.353, z=0.927, ranksum=11134). (Note that the difference between both and Simon survives Bonferroni correction). Despite the non-significant difference between both conflict trials and Eriksen-only trials, when the effects of either single type of conflict (Eriksen or Simon) are averaged, the effect of both types of conflict occurring together is still larger than the effect of either one (p=0.012, z=2.514, ranksum=15093).

### Pre-trial single neuron correlates of conflict

We recorded from 378 neurons from 9 patients in dlPFC. Our goal was to determine whether responses of neurons before the start of the trial predict subsequent reaction time. Consider, first, responses of an example neurons shown in **Figure 2**.

**Figure 2.**
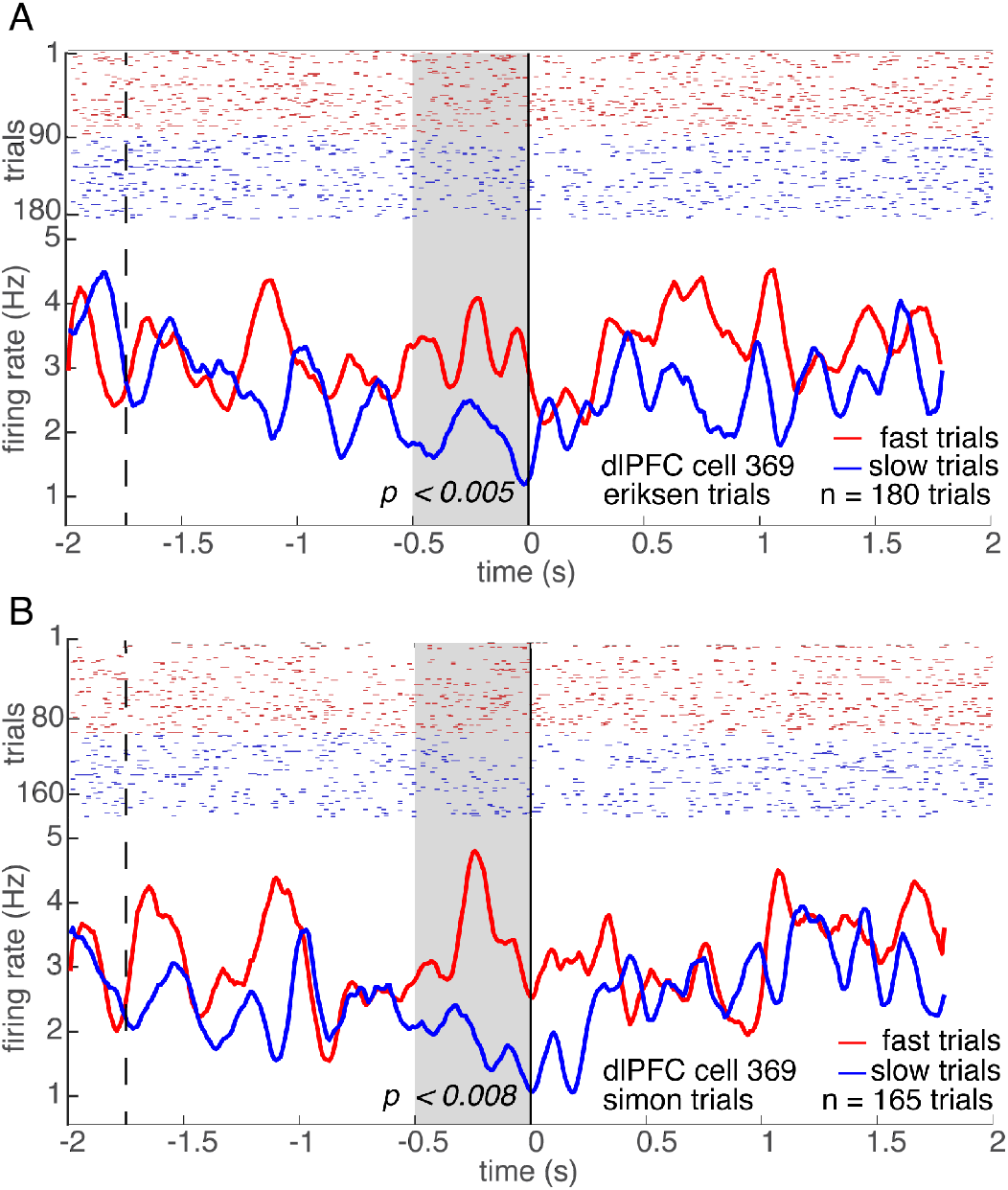
Individual dlPFC neurons signal the speed of upcoming responses in a conflict-specific manner. (A-B) PSTHs of example neuron 369 showing significantly higher pre-trial activity before fast (red) responses than slow (blue) responses in both (A) Eriksen and (B) Simon conflict trials. Gray shading indicates the 500ms analysis window. The dotted line indicates the response on the last trial and the solid vertical line indicates the stimulus onset.

**Figure 3.**
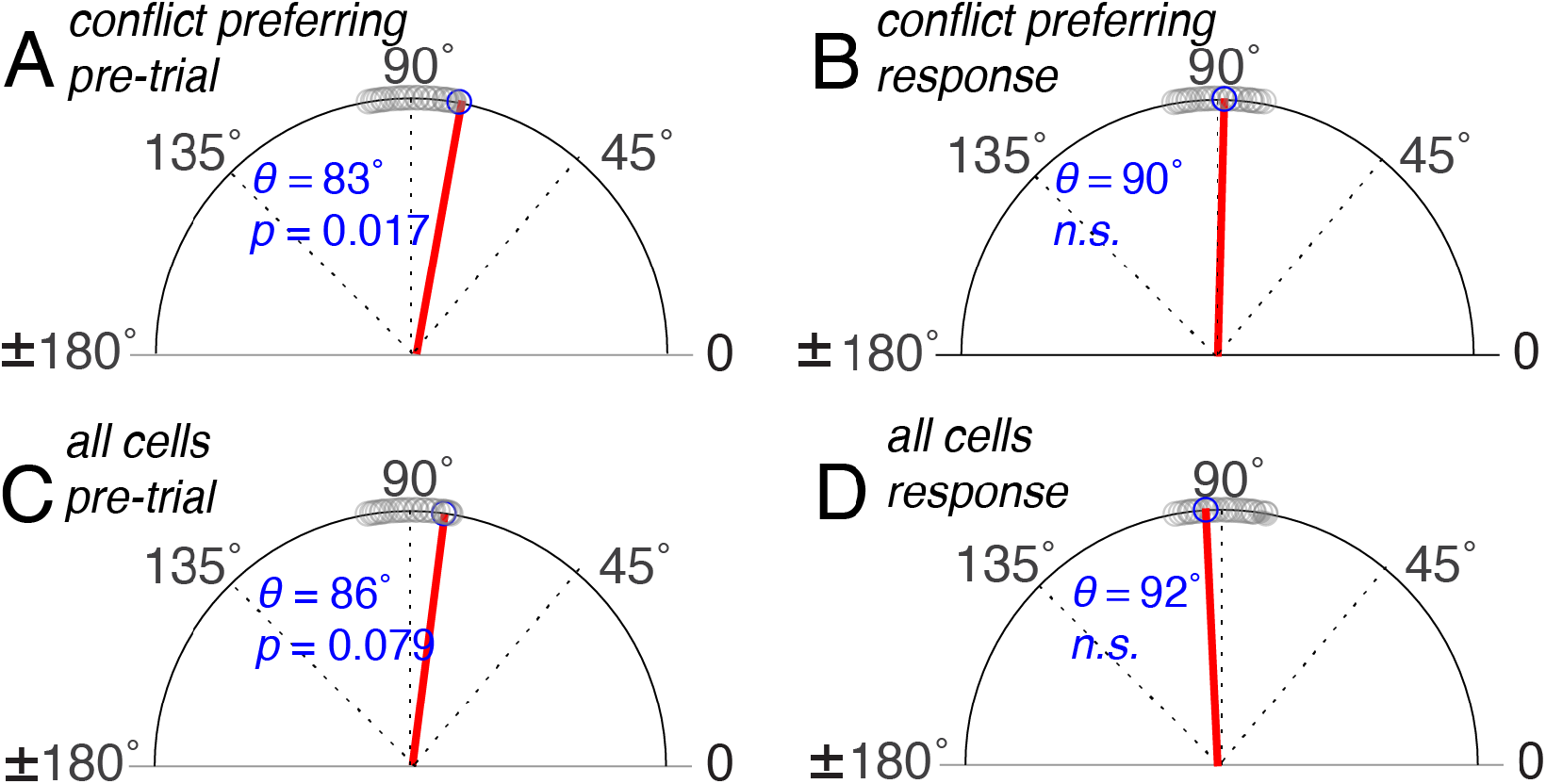
The angle between conflict-response tuning weight vectors in dlPFC orthogonalizes between the pre-trial and response period. A zero-degree angle represents complete collinearity. (A-D) the angle between Eriksen and Simon coding vectors (blue circle, blue numbering and red line). The significance of the difference from a 90° (blue italics) was computed from permutation testing. The null distribution is shown with grey circles. (A) Vector angle between conflict preferring cells (amodal or conflict-type specific) showing superimposed collinear and orthogonal coding. (B) The angle between all cells shows a similar trend. (C) Same as (A) for response codes, showing purely orthogonal coding. (D) Same as for (B) for response codes, showing orthogonal coding.

**Figure 4.**
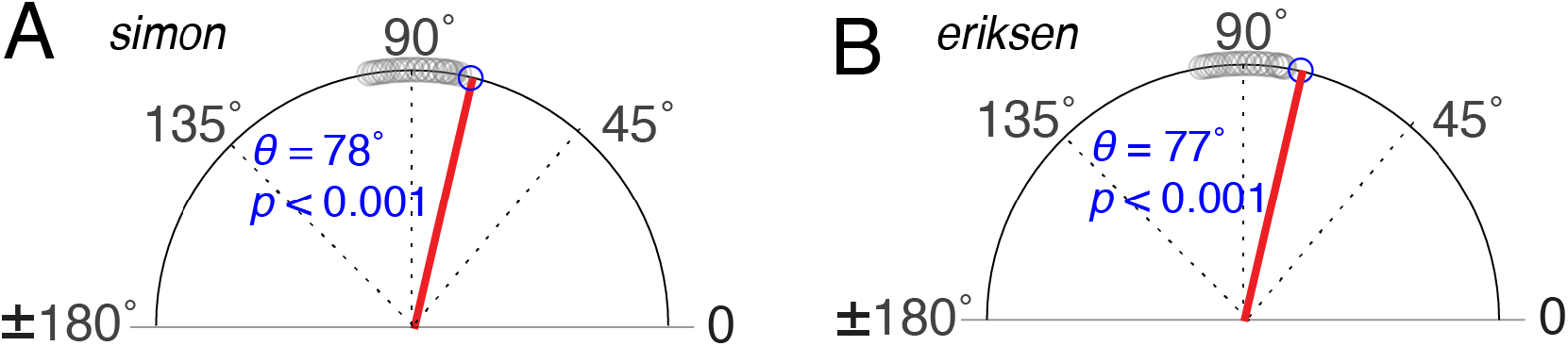
The angle between pre-trial and response conflict-response tuning weight vectors in dlPFC shows partially co-linear coding in dlPFC. A zero-degree angle represents complete collinearity. (A-B) Conflict-preferring coding vector angles (blue circle, blue numbering and red line); the significance of the difference from a 90° (blue italics) was computed from permutation testing (the null distribution is shown with grey circles). (A) the angle between pre-trial and response Eriksen coding vectors shows significant collinearity as does (B) the angle between pre-trial and response Simon coding vectors.

In our example neuron (**Figure 2**), taken from dlPFC, both Simon responses and Eriksen responses were significantly greater before the start of faster reaction time trials than before slower reaction time trials (fast Simon RT trials: 3.2 spikes/sec, slow Simon RT trials: 2.5 spikes/sec, p=0.005, t-test on z-scored data; fast Eriksen RT trials: 3.2 spikes/sec, slow Eriksen RT trials: 2.5 spikes/sec, p=0.008, t-test on z-scored data). Specifically, we ran a median split on reaction times post-hoc and separated firing rates on those two categories. As described below, firing rate in this neuron also predicted the reaction time in a continuous model.

### Pre-trial population correlates of conflict

To explore these effects at the population level, we fit generalized linear models (GLMs) to the pretrial firing rates and reaction times for all trials. Our analyses controlled for prior trial conflict type, because conflict level on the previous trial can modulate preparatory neural responses and lead to trial-to-trial adjustments, such as post-error slowing and conflict-adaptation/trial congruency effects ^19,31,32^. Our analyses also controlled for previous trial reaction time (RT), to remove possible effects of slow drifts in arousal. We analyzed the 500ms epoch between the fixation and cue onset in the current trial.

We first asked whether pre-trial activity predicted reaction times on all trials without regard to conflict or history. Pre-trial firing rates in 13.2% of neurons in dlPFC were predictive of the RT on the upcoming trial (*n* = 50/378 neurons). This proportion is greater than that predicted by chance (*p* < 0.001, one-sided binomial test) and remained the same after controlling for prior trial reaction time and conflict type. These results indicate that a small but statistically significant fraction of neurons have firing rates that predict upcoming reaction times.

After establishing that pretrial activity predicts reaction time overall, we next examined whether it modulated reaction times differently depending on the upcoming conflict condition. We compared a model with a single parameter for any conflict type (“conflict type-amodal”, valued at 1 for any conflict type and 0 for no conflict) to a model with separate parameters for Eriksen and Simon conflict types (“conflict type-specific”), including trials in which the distractor stimuli appeared separately as well as together, and controlling for previous trial conflict and RT. We also compared these models to the “no-conflict” model described above. In both areas, we found significant proportions of cells with a significant main effect of firing rate on response time.

At the single cell level, we found evidence for both conflict type-amodal tuning and conflict type-specific tuning. In the conflict type-amodal model, 7.7% of cells exhibited significant single conflict coefficients, a proportion that was significantly greater than chance (*n* = 29/378, *p* = 0.009, binomial test with chance rate of 5%) and 11.1% of cells showed a main effect (Wald test) of firing rate (*p* < 0.001, one sided binomial test with chance rate of 5%). With the conflict type-specific model, 14.6 % of all cells (*n*=55/378) exhibited significant coefficients (Wald tests) for predicting RT on either Eriksen trials or on Simon Trials (*p* < 0.001, binomial test with chance rate of 9.75%). Examining the main effect of overall firing rate in this model, 13.2% of cells showed a significant effect (*n* = 50/378,*p* < 0.001, binomial test).

When we examined the evidence for conflict tuning in dlPFC with model comparison, we found that 73% of cells preferred the conflict-specific model (*n* = 277/378, median model 3:1 BIC weight ratio = 23, median model 3:2 BIC weight ratio = 40), 22% preferred the no-conflict model (*n* = 83/378, median model 1:3 BIC weight ratio = 2.9, median model 1:2 BIC weight ratio = 44) and only 3% preferred the conflict type-amodal model (*n* = 10/378, median model 2:3 BIC weight ratio = 2.8, median model 2:1 BIC weight ratio = 2.4 x 10^6^). Of the 29 cells with significant conflict type-amodal coefficients, 93% (*n* = 27/29) preferred the conflict type-specific model. These BIC results indicate that cells tuned for conflict type-specific responding dominate in dlPFC.

### Semi-orthogonal coding for two forms of conflict preparation in dlPFC

Our observation that a significant portion of individual cells code for conflict type-amodal reaction times but yet were mostly better described with the conflict type-specific model led us to wonder if *population-level* activity might contain a conflict type-amodal signal. To test this possibility, we asked how ensemble codes for predicting resolution of Simon and Eriksen conflict, derived from the conflict type-specific model, were related to each other. To do this, we calculated a vector of GLM regression weights for each distractor type across neurons. We call these vectors the *pre-trial tuning weight vectors*. We then compared these vectors by computing the Spearman correlation of the vectors corresponding to Eriksen and Simon coefficients (cf. ^33^). (Note that the Spearman test does not assume linearity and is thus more general than the more common Pearson and is also less sensitive to potential outliers). We tested these analyses on all cells that preferred either conflict type-amodal or conflict type-specific models by BIC (*n* = 287/738) as well as on all cells; doing so allows us to include contributions from all relevant neurons, even those with real effects that do not pass the strict significance threshold; this approach thus has better signal-to-noise (and moderately less susceptible to Type II errors) than analysis approaches that focus on cells that cross a significance threshold (Blanchard et al., 2018).

We found that in the conflict-preferring population, the codes for Simon and Eriksen were weakly positively correlated (*ρ* = 0.12, *p* = 0.048; permutation test). Relatedly, we found the angle *θ* between these vectors to be 83° and significantly less than 90° (*p* = 0.017, permutation test). This result is consistent with the population-level superposition of collinear and orthogonal coding for optimal initial conditions for conflict responding. In the entire population, the angle *θ* between these vectors was 86° and less than 90° at a trend level (*p* = 0.079, permutation test), and the Spearman’s *ρ* was 0.093 (*p* = 0.038, permutation test). In other words, this result indicates that there is a conflict type-amodal code recoverable from dlPFC ensemble activity.

### Firing rates in the response period explain response time variability

If neural firing patterns in the preparatory period bias response speed in a conflict type-specific manner, we would expect the neural state in the response period to reflect this. To test this hypothesis, we ran the conflict type-specific model described above on the 500ms period after stimulus onset. We found that 16.9 % of all cells (*n* = 64/378) exhibited significant coefficients (Wald Test) for predicting RT on either Eriksen trials or Simon Trials (*p* < 0.001, binomial test with chance rate of 9.75%). To examine the degree to which the preparatory state resembled the response state, we computed similarity measures for their respective coding vectors, which we call the *response tuning weight vectors*. For both areas, we found that the pre-trial and response tuning weight vectors were partially co-linear. The Spearman correlation between Eriksen codes for conflict-preferring cells was r = 0.22 (*p* < 0.001, permutation test) and between Simon codes was *ρ* = 0.23 (*p* < 0.001, permutation test); the angles were 77° for the Eriksen (*p* < 0.001, permutation test) and 78° for the Simon condition (*p* < 0.001, permutation test). The results for all cells were similar (Eriksen *ρ* = 0.22, *p* < 0.001 and *θ* = 78°, *p* < 0.001; Simon *ρ* = 0.23, *p* < 0.001 and *θ* = 75°, *p* < 0.001). These results support the notion that the neural states in dlPFC that predispose for more efficient conflict resolution overlap with the states of efficient responding themselves. Next, we queried the association between the Eriksen and Simon response tuning weight vectors. We found that the codes for both the conflict-preferring and all-cell populations were fully orthogonalized in the response period, in contrast to the pre-trial period (conflict-preferring cells *ρ* = −0.01, p = 0.882 and *θ* = 90°, *p* > 0.05; all cells *ρ* = −0.04, *p* > 0.05 and *θ* = 92°, *p* > 0.05, permutation tests). To determine whether the pre-trial and response correlation coefficients and angles differed, we computed bootstrap distributions for each and compared the medians. The correlation coefficient for Eriksen versus Simon pre-trial tuning weight vectors differed significantly from those for the response tuning weight vectors for both the conflict-preferring population and all cells (Wilcoxon rank sums, all *p* <0.001). This is important to confirm because the difference between a significant and non-significant effect is not necessarily itself significant ^34^.

## DISCUSSION

We examined the responses of single neurons in human dlPFC during a task that interleaved two kinds of conflict, Simon and Eriksen. We found that firing rates of a modest but significant proportion of neurons in both areas before the trial predicts the efficiency of cognitive control, as inferred from reaction times. The fact that ensemble responses predict reaction time before the trial, and presumably task-driven cognition, supports the hypothesis that successful cognitive control reflects, in part, the ability to transition through specific brain states. By controlling for prior trial condition and reaction time, we showed that these brains states do not simply reflect adaptation or drifts in arousal, but rather history-independent patterns that predispose to efficient responding.

We also compared the patterns that predicted the efficiency of upcoming responses on trials with either Simon, Eriksen or both types of conflict. At the individual cell level, we found that these responses were consistent with three types of codes: a conflict-independent (no-conflict), a conflict type-amodal and conflict type-specific code. At the population level, we found a weak general conflict type-amodal code. These results demonstrate, first, that neurons in dlPFC process both type-specific and domain-general neural computations. While the domain-general element was observed both at the individual cell level and the population level, model comparison suggests it is more likely a population level phenomenon: in most cells the full model with separate conflict terms significantly outperformed a model with one single term for any type of conflict. A sizable proportion of individual cells (22%) did prefer a model with no conflict term, predicting response speed across all conditions. However, the domain-general/conflict type-amodal element emerged at the population level from the correlation between the conflict-specific regression coefficients across neurons.

Studies of motor control in rodents and non-human primates have shown that variability in the firing rates of neurons in motor cortex during movement preparation predicts variability in subsequent movements ^6,9,10^. This body of work has given rise to the “initial condition hypothesis”, which posits that neural firing patterns behave like dynamical systems, and the trajectory of neural dynamics therefore depends on the initial state of the system ^3,8,35^. One implication of this theory is that preparatory neural states can therefore be optimized to efficiently produce a desired behavioral outcome ^36^. If similar principles apply to neural dynamics during cognitive tasks, this suggests that preparatory activity may also be optimized to support the efficient application of cognitive control, consistent with the notion of pro-active control ^37^.

Indeed, a growing literature suggests neural patterns subserving cognition are also consistent with dynamical systems models ^38^. Using the same task studied, here, we recently showed that neural population activity in prefrontal regions during conflict resolution follows low-dimensional trajectories that differ depending on the type of conflict ^28^. The divergence of these trajectories raises the question of whether they have different optimal initial states, or if similar neural starting conditions give rise to similar task performance. The results we present provide a somewhat nuanced answer: the optimal initial states overlap slightly, suggesting a weak mechanism for shared conflict-responding, but orthogonalize over the course of response, supporting the notion of trajectory divergence.

What are the implications of preparatory brain states for conflict resolution? One possibility is that initial states that predict response times reflect changes in arousal or attention that are either spontaneous or the result of adaptations to the previous trial. While attractive for its simplicity, this explanation would not explain the persistence of the effect after controlling for trial history nor would it explain the conflict type-specific nature of the initial states. Rather, our results suggest that the computations that perform conflict resolution are not only distinct, as implied by the divergent neural state space trajectories, but are facilitated with both shared and independent factors. In this task, participants do not have knowledge of whether the upcoming trial will have one form of conflict or the other or both. Maintaining both shared and independent preparatory factors for conflict types, independent of the past, may allow the brain to respond flexibly to unpredictable challenges while minimizing interference between the processes needed to respond to those challenges. Such a factorized model of representation has been proposed as a mechanism to minimize interference and facilitate generalization in learning ^39^. Why would preparatory states vary from trial to trial? The simplest explanation is spontaneous fluctuations due to noise. Another possibility is that participants are subtly making predictions about the condition of the upcoming trial, perhaps based on longer trial history than the recent past controlled for here.

Recent years have seen the emergence of the dynamical systems perspective in motor neurophysiology. This view sees the aggregate activity of neurons in a region as constituting a state and that the execution of motor actions is driven by the lawful progression across states in motor regions. It has been further proposed that cognitive performance may reflect a similar dynamical system view, however, this view has been difficult to test. We recently demonstrated that, in asynchronous choice, neurons in two core reward areas show subspace orthogonalization, a neural process previously associated only with motor cortex ^40^. Here we sought to test a critical prediction of the initial state hypothesis. In the motor cortex, the *initial state hypothesis* holds that successful implementation of a motor actions cannot begin until motor cortex enters into a specific ensemble state, defined in practice as an ensemble firing rate pattern. We hypothesized that an analogous idea in cognition would be that implementation of a cognitive act would require implementation of an initial state. Our results suggest that not only does the initial state of neural activity prior to the cognitive act of conflict resolution support efficient responding in a pro-active manner ^37^, it does so in a largely conflict type-specific manner. The small degree of collinearity we observe in dlPFC may contribute to untangling of stimulus-action processes before full orthogonalization during implementation. Previous work in premotor cortex has shown that the neural state space during responding predicts reaction times in motor tasks ^41^. We found that these optimal initial states share structure with the optimal responding states, suggesting they support optimal response trajectories. Future studies may examine whether neural state-space trajectories separate by response time differently depending on both the initial states and the specific cognitive computations the brain performs.

## METHODS

### Subjects and ethics statement

We studied one cohort of 9 patients: 8 (2 female) with movement disorders (Parkinson’s disease or essential tremor) who were undergoing deep brain stimulation (DBS) surgery, and one male patient with epilepsy undergoing intracranial seizure monitoring. The entry point for the trajectory of the DBS electrode is typically in the inferior portion of the superior frontal gyrus or superior portion of the middle frontal gyrus, within 2 cm of the coronal suture. This area corresponds to dlPFC (Brodmann’s areas 9 and 46). The single epilepsy patient in this cohort underwent a craniotomy for placement of subdural grid/strip electrodes in a prefrontal area including dlPFC.

All decisions regarding sEEG and DBS trajectories and craniotomy location were made solely based on clinical criteria. The Columbia University Medical Center Institutional Review Board approved these experiments, and all subjects provided informed consent prior to participating in the study.

### Behavioral Task

All subjects performed the multi-source interference task (MSIT; **Figure 1A**). In this task, each trial began with a 500-millisecond fixation period. This was followed by a cue indicating the ***correct response*** as well as the ***distractor response***. The cue consisted of three integers drawn from {0, 1, 2, 3}. One of these three numbers (the “***correct response cue***”) was different from the other two numbers (the “***distractor response cues***”). Subjects were instructed to indicate the identity of the correct response number on a 3-button pad. The three buttons on this pad corresponded to the numbers 1 (left button), 2 (middle) and 3 (right), respectively.

The MSIT task therefore presented two types of conflict. Simon (motor spatial) conflict occurred if the correct response cue was located in a different position in the cue than the corresponding position on the 3-button pad (e.g. ‘0 0 1’; target in right position, but left button is correct choice). Eriksen (flanker) conflict occurred if the distractor numbers were possible button choices (e.g. ‘3 2 3’, in which “3” corresponds to a possible button choice; vs. ‘0 2 0’, in which “0” does not correspond to a possible button choice).

After each subject registered his or her response, the cue disappeared, and feedback appeared. The feedback consisted of the target number, but it appeared in a different color. The duration of the feedback was variable (300 to 800 milliseconds, drawn from a uniform distribution therein). The inter-trial interval varied uniformly randomly between 1 and 1.5 seconds.

The task was presented on a computer monitor controlled by the Psychophysics Matlab Toolbox (www.psychtoolbox.org; The MathWorks, Inc). This software interfaced with data acquisition cards (National Instruments,) that allowed for synchronization of behavioral events and neural data with sub-millisecond precision.

### Data Acquisition and preprocessing

Single unit activity (SUA) was recorded from a combination of two techniques. The DBS surgeries were performed according to standard clinical procedure, using clinical microelectrode recording (Frederick Haer Corp.). Prior to inserting the guide tubes for the clinical recordings, we placed the microelectrodes in the cortex under direct vision to record from dlPFC, (IRB-AAAK2104). The epilepsy implant in Cohort 2 included a Utah-style microelectrode array (UMA) implanted in dlPFC (IRB-AAAB6324). In all cases, data were amplified, high-pass filtered, and digitized at 30 kilosamples per second on a neural signal processor (Blackrock Microsystems, LLC).

SUA data were re-thresholded offline at negative four times the root mean square of the 250 Hz high-pass filtered signal. Well-isolated action potential waveforms were then segregated in a semi-supervised manner using the T-distribution expectation-maximization method on a feature space comprised of the first three principal components using Offline Sorter (OLS) software (Plexon Inc, Dallas, TX; USA). The times of threshold crossing for identified single units were retained for further analysis.

### Data Analysis

We determined the effect variations in pre-stimulus firing rates had on reaction times by comparing four generalized linear models. We first fit gamma distributions to the reaction times and excluded reaction times with a less than 0.005 probability following ^42^. For each model, we centered and scaled the continuous predictor variables (firing rate and reaction time) by z-scoring. We analyzed correct and incorrect trials to prevent false-positives from data-censoring effects. We pre-selected the pre-trial analysis interval as the 500ms period between fixation and stimulus onset and the response analysis interval as the 500ms following stimulus onset. To determine the overall effect of firing rate marginalized over condition, we first fit the following generalized linear model:

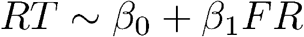

We then compared several alternative models, while controlling for reaction time and prior conflict: a single firing rate coefficient for all trials model,

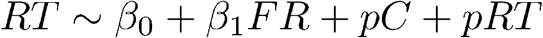

a model with an additional term for the firing rate on trials with any conflict (Simon, Eriksen or both),

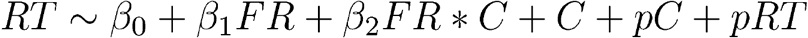

and a model with separate, additive conflict terms for Simon and Eriksen conditions,

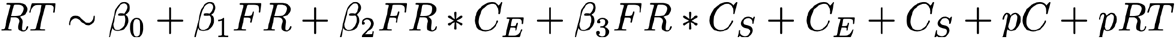

where FR is firing rate on all trials, C is an indicator variable for trials with any conflict, CE and CS are indicator variables for Eriksen and Simon trials, respectively, pC is a categorical variable for prior conflict type and pRT is the previous trial reaction time. We used a normal distribution with a log link function because the reaction time data was well described by a log-normal distribution.

We compared these models by their BIC weight ratios ^43^. We identified the best fitting model for each cell as the model with a greater than one BIC weight ratio for all pairwise comparisons. The BIC penalty for model complexity is greater than that for Akaike Information Criteria (AIC) as the number of model parameters exceeds e^2^ ~ 7 and thus more appropriate here (and more conservative) ^43,44^.To assess the significance of overall model fits we performed deviance tests relative to a constant model. To assess the significance of coefficients we performed Wald tests (REF).

To compare the neural codes for conflict, we entered the regression coefficients into individual vectors for each conflict condition. We then computed the Spearman correlation between those vectors as well as the angle between the vectors. We excluded points more than 3 median absolute deviations from the median because angle measurements; this approach excluded 6 cells. We then randomized the vector entries and computed correlations between the randomized vectors to form null distributions (2000 permutations). We computed p-values for the real measurement relative to the corresponding null distribution (permutation test). To compare angles and correlations, we constructed bootstrap distributions with resampling (5000 samples) and compared the medians of the resultant distributions with the non-parametric Wilcoxon rank sum test.

## Acknowledgements

This work was supported by NIH R01 MH106700, NIH U01 NS108923, NIH K12 NS080223, NIH S10 OD018211, NIH R01 NS084142, NIH R01 DA038615, the Brain and Behavior Foundation, and the Dana Foundation. This work was also supported by the Center for Neuroengineering (CNE) at UMN. Special thanks to Camilla Casadei, David K. Peprah, and Timothy G. Dyster for coordination and data collection efforts. The funders had no role in study design, data collection and analysis, decision to publish, or preparation of the manuscript.

## References

1. Weissman, D. H., Roberts, K. C., Visscher, K. M. & Woldorff, M. G. The neural bases of momentary lapses in attention. Nat. Neurosci. 9, 971–978 (2006).

2. Boly, M. et al. Baseline brain activity fluctuations predict somatosensory perception in humans. Proc. Natl. Acad. Sci. U. S. A. 104, 12187–12192 (2007).

3. Churchland, M. M. et al. Stimulus onset quenches neural variability: a widespread cortical phenomenon. Nat. Neurosci. 13, 369–378 (2010).

4. Pezzulo, G. & Cisek, P. Navigating the Affordance Landscape: Feedback Control as a Process Model of Behavior and Cognition. Trends Cogn. Sci. 20, 414–424 (2016).

5. Riehle, A. & Requin, J. Monkey primary motor and premotor cortex: single-cell activity related to prior information about direction and extent of an intended movement. J. Neurophysiol. 61, 534–549 (1989).

6. Churchland, M. M., Yu, B. M., Ryu, S. I., Santhanam, G. & Shenoy, K. V. Neural variability in premotor cortex provides a signature of motor preparation. J. Neurosci. 26, 3697–3712 (2006).

7. Churchland, M. M., Cunningham, J. P., Kaufman, M. T., Ryu, S. I. & Shenoy, K. V. Cortical preparatory activity: representation of movement or first cog in a dynamical machine? Neuron 68, 387–400 (2010).

8. Shenoy, K. V., Sahani, M. & Churchland, M. M. Cortical control of arm movements: a dynamical systems perspective. Annu. Rev. Neurosci. 36, 337–359 (2013).

9. Afshar, A. et al. Single-trial neural correlates of arm movement preparation. Neuron 71, 555–564 (2011).

10. Pandarinath, C. et al. Inferring single-trial neural population dynamics using sequential auto-encoders. Nat. Methods 15, 805–815 (2018).

11. Becket Ebitz, R. & Hayden, B. Y. The population doctrine revolution in cognitive neurophysiology. arXiv [q-bio.NC] (2021).

12. Churchland, A. K., Kiani, R. & Shadlen, M. N. Decision-making with multiple alternatives. Nat. Neurosci. 11, 693–702 (2008).

13. Pastor-Bernier, A. & Cisek, P. Neural Correlates of Biased Competition in Premotor Cortex. Journal of Neuroscience vol. 31 7083–7088 (2011).

14. Yoo, S. B. M. & Hayden, B. Y. Economic Choice as an Untangling of Options into Actions. Neuron 99, 434–447 (2018).

15. Hunt, L. T. & Hayden, B. Y. A distributed, hierarchical and recurrent framework for reward-based choice. Nat. Rev. Neurosci. 18, 172–182 (2017).

16. Smith, E. H. et al. Widespread temporal coding of cognitive control in the human prefrontal cortex. Nat. Neurosci. 22, 1883–1891 (2019).

17. Ebitz, R. B. & Platt, M. L. Neuronal activity in primate dorsal anterior cingulate cortex signals task conflict and predicts adjustments in pupil-linked arousal. Neuron 85, 628–640 (2015).

18. Bryden, D. W. et al. Single Neurons in Anterior Cingulate Cortex Signal the Need to Change Action During Performance of a Stop-change Task that Induces Response Competition. Cereb. Cortex 29, 1020–1031 (2019).

19. Sheth, S. A. et al. Human dorsal anterior cingulate cortex neurons mediate ongoing behavioural adaptation. Nature 488, 218–221 (2012).

20. Shenhav, A. et al. Toward a Rational and Mechanistic Account of Mental Effort. Annu. Rev. Neurosci. 40, 99–124 (2017).

21. Yeung, N., Holroyd, C. B. & Cohen, J. D. ERP correlates of feedback and reward processing in the presence and absence of response choice. Cereb. Cortex 15, 535–544 (2005).

22. Botvinick, M. M., Braver, T. S., Barch, D. M., Carter, C. S. & Cohen, J. D. Conflict monitoring and cognitive control. Psychol. Rev. 108, 624–652 (2001).

23. Botvinick, M. M. & Cohen, J. D. The Computational and Neural Basis of Cognitive Control: Charted Territory and New Frontiers. Cognitive Science vol. 38 1249–1285 2014).

24. Miller, E. K. & Cohen, J. D. An integrative theory of prefrontal cortex function. Annu. Rev. Neurosci. 24, 167–202 (2001).

25. MacDonald, A. W., 3rd, Cohen, J. D., Stenger, V. A. & Carter, C. S. Dissociating the role of the dorsolateral prefrontal and anterior cingulate cortex in cognitive control. Science 288, 1835–1838 (2000).

26. Shenhav, A., Botvinick, M. M. & Cohen, J. D. The expected value of control: an integrative theory of anterior cingulate cortex function. Neuron 79, 217–240 (2013).

27. Johnston, K., Levin, H. M., Koval, M. J. & Everling, S. Top-down control-signal dynamics in anterior cingulate and prefrontal cortex neurons following task switching. Neuron 53, 453–462 (2007).

28. Becket Ebitz, R. et al. Human dorsal anterior cingulate neurons signal conflict by amplifying task-relevant information. Cold Spring Harbor Laboratory 2020.03.14.991745 (2020) doi:10.1101/2020.03.14.991745.

29. Bush, G., Shin, L. M., Holmes, J., Rosen, B. R. & Vogt, B. A. The Multi-Source Interference Task: validation study with fMRI in individual subjects. Mol. Psychiatry 8, 60–70 (2003).

30. Deng, Y., Wang, X., Wang, Y. & Zhou, C. Neural correlates of interference resolution in the multi-source interference task: a meta-analysis of functional neuroimaging studies. Behav. Brain Funct. 14, 8 (2018).

31. Horga, G. et al. Adaptation to conflict via context-driven anticipatory signals in the dorsomedial prefrontal cortex. J. Neurosci. 31, 16208–16216 (2011).

32. Oehrn, C. R. et al. Neural communication patterns underlying conflict detection, resolution, and adaptation. J. Neurosci. 34, 10438–10452 (2014).

33. Blanchard, T. C., Hayden, B. Y. & Bromberg-Martin, E. S. Orbitofrontal cortex uses distinct codes for different choice attributes in decisions motivated by curiosity. Neuron 85, 602–614 (2015).

34. Nieuwenhuis, S., Forstmann, B. U. & Wagenmakers, E.-J. Erroneous analyses of interactions in neuroscience: a problem of significance. Nat. Neurosci. 14, 1105–1107 (2011).

35. Ames, K. C. & Churchland, M. M. Motor cortex signals for each arm are mixed across hemispheres and neurons yet partitioned within the population response. Elife 8, (2019).

36. Kao, C.-H., Lee, S., Gold, J. I. & Kable, J. W. Neural encoding of task-dependent errors during adaptive learning. Elife 9, (2020).

37. Braver, T. S. The variable nature of cognitive control: a dual mechanisms framework. Trends Cogn. Sci. 16, 106–113 (2012).

38. Taghia, J. et al. Uncovering hidden brain state dynamics that regulate performance and decision-making during cognition. Nature Communications vol. 9 (2018).

39. Behrens, T. E. J. et al. What Is a Cognitive Map? Organizing Knowledge for Flexible Behavior. Neuron 100, 490–509 (2018).

40. Yoo, S. B. M. & Hayden, B. Y. The Transition from Evaluation to Selection Involves Neural Subspace Reorganization in Core Reward Regions. Neuron 105, 712–724.e4 (2020).

41. Michaels, J. A., Dann, B., Intveld, R. W. & Scherberger, H. Predicting Reaction Time from the Neural State Space of the Premotor and Parietal Grasping Network. J. Neurosci. 35, 11415–11432 (2015).

42. Widge, A. S. et al. Deep brain stimulation of the internal capsule enhances human cognitive control and prefrontal cortex function. Nat. Commun. 10, 1536 (2019).

43. Wagenmakers, E.-J. & Farrell, S. AIC model selection using Akaike weights. Psychon. Bull. Rev. 11, 192–196 (2004).

44. Model Selection and Multimodel Inference: A Practical Information-Theoretic Approach. (Springer, New York, NY, 2002).

